# Polyelectrolyte Coating of Ferumoxytol Differentially Impacts on Labeling of Inflammatory and Steady-State Dendritic Cell Subtypes

**DOI:** 10.1101/2021.05.27.445994

**Authors:** Nehar Celikkin, John E. Wong, Martin Zenke, Thomas Hieronymus

## Abstract

Engineered magnetic nanoparticles (MNPs) are emerging as advanced tools for medical applications. The coating of MNPs using polyelectrolytes (PEs) is a versatile means to tailor MNP properties and is used to optimize MNP functionality. Dendritic cells (DCs) are critical regulators of adaptive immune responses. Functional distinct DC subsets exist either under steady-state or inflammatory conditions, which are explored for the specific treatment of various diseases, such as cancer, autoimmunity or transplant rejection. Here, the impact of PE coating of ferumoxytol for uptake into both inflammatory and steady-state DCs and cellular responses to the MNP labeling is addressed. Labeling efficiency by uncoated and PE-coated ferumoxytol is highly variable in different DC subsets, and PE coating significantly improves the labeling of steady-state DCs. Uncoated ferumoxytol results in increased cytotoxicity of steady-state DCs after labeling that is abolished by the PE coating, while no increased cell death is observed in inflammatory DCs. Furthermore, uncoated and PE-coated ferumoxytol appears immunologically inert in inflammatory DCs but induces activation of steady-state DCs. These results show that PE coating of MNPs can be applied to endow particles with desired properties for enhanced uptake and cell type-specific responses in distinct target DC populations.

## Introduction

Cellular immunotherapies become an increasingly promising approach for the development of integrative and personalized therapies. Antigen-presenting dendritic cells (DCs) are considered as particularly well suited for the development of such therapies due to their unique capacity to initiate and orchestrate antigen-specific immune responses [1]. DCs are the first cells to be involved in antigen sensing and scavenging, followed by processing and subsequent antigen-specific T-cell priming. Additionally, DCs can activate further immune effector cells, including B cells, natural killer (NK) cells and NKT cells [2]. Thus, DCs harmonize immune responses that eventually result in resistance to foreign pathogens and tolerance to self. Novel DC-based therapies, therefore, aim at establishing beneficial immune conditions in diseases such as cancer, chronic inflammation, autoimmunity, or transplant rejection. This is readily achieved using DC-targeted vaccines, mainly to induce anti-tumor immunity, while tolerogenic DCs are explored to silence autotoxic immune responses [3–6].

*In vivo* monitoring of engraftment, position and/or migration to the target site and function of transplanted cells could decisively contribute to the success of such therapies. Magnetic resonance imaging (MRI) of contrast agent-labeled cells, including DCs, has emerged as a well-suited imaging technique for tracking cells *in vivo* [7]. An outstanding feature of MRI is its capacity for long-term tracking of cells and their and migration while achieving excellent high-resolution images of target tissue in a three-dimensional anatomical context. Stable labeling of DCs with iron oxide-based magnetic nanoparticles (MNPs) as contrast agents has proven successful for MRI-based detection of cell deposits and their migration [8–12].

Ferumoxytol is an FDA approved iron oxide-based MNP formulation currently used as a drug to treat iron-deficiency anemia in chronic kidney disease [13]. Ferumoxytol has also been used as a macrophage-imaging agent as well as a blood-pool agent with MRI [14]. However, ferumoxytol alone did not result in effective cell labeling [15,16]. Thus, in previous studies, we successfully established conditions that enabled us to manufacture colloidal stable ferumoxytol particles coated with polyelectrolytes (PE), resulting in enhanced cell labeling [16]. In the present study, we mainly focused on the impact of PE-coated ferumoxytol MNPs on the labeling of different DC subsets, including inflammatory DCs, and steady-state conventional DCs (cDCs) and plasmacytoid DCs (pDCs) [17]. DCs generated *in vitro* from monocytes or CD34+ hematopoietic stem/progenitor cells (HSC) of blood or bone marrow using the granulocyte-macrophage colony-stimulating factor (GM-CSF) resemble a DC subset that occurs *in vivo* only under inflammatory conditions and thus referred to as inflammatory DCs [17]. Such patient-derived DC represents the prevailing DC subtype used in autologous cell-based immunotherapies so far [6]. However, employing primary existing cDCs and pDCs subtypes under steady-state conditions for therapeutic approaches are considered as potentially better-suited alternatives [3]. Here, we investigated the cellular intake of PE-coated and unmodified ferumoxytol into inflammatory DCs, and steady-state cDCs and pDCs generated *in vitro* cultures from mouse bone marrow. We assessed the impact of the PE coatings on cell viability, labeling efficiency, and intracellular iron content of MNP-labeled cells. Furthermore, immunophenotypic alterations of the various DC subtypes upon MNP incorporation were addressed. In summary, the results reported here revealed a differential impact of PE-coated and uncoated MNPs upon uptake in inflammatory DCs and steady-state DC, respectively.

## Materials and Methods

### Polyelectrolyte coating of ferumoxytol

The FDA approved iron oxide nanoparticle formulation ferumoxytol (Rienso®) was purchased from Takeda Pharma (Konstanz, Germany). High MW polyethyleneimine (PEI, 750 kDa) and low MW polydiallyldimethylammoniumchlorid (PDADMAC, 100-200 kDa) were obtained from Sigma-Aldrich (Taufkirchen, Germany). The surface coating of ferumoxytol was carried out using the LbL technique to deposit polyelectrolyte layers (Figure 1 A) [16,18–20]. The negatively charged ferumoxytol MNPs were added to an aqueous solution of the positively charged polyelectrolytes to prepare the coating layer. For coating with PEI, 3 mg of ferumoxytol MNPs were added into 5 mL of a 1g/L PEI solution. For coating with PDADMAC, 9 mg of ferumoxytol MNPs were added into 5 mL of a 2g/L PDADMAC solution [16]. After overnight shaking, the products were separated from the excess polyelectrolyte by magnetic separation and rinsed 3 times with double distilled water (ddH2O). Hydrodynamic diameter (dH) of MNPs was obtained from dynamic light scattering (DLS) by cumulant fits using a Zetasizer 3000HSA (Malvern Instruments, Malvern, UK), which also provides the zeta potentials of the particles; each value reported is an average of at least ten consistent measurements. Uncoated and coated MNPs were visualized by transmission electron microscopy (TEM).

**Figure 1.**
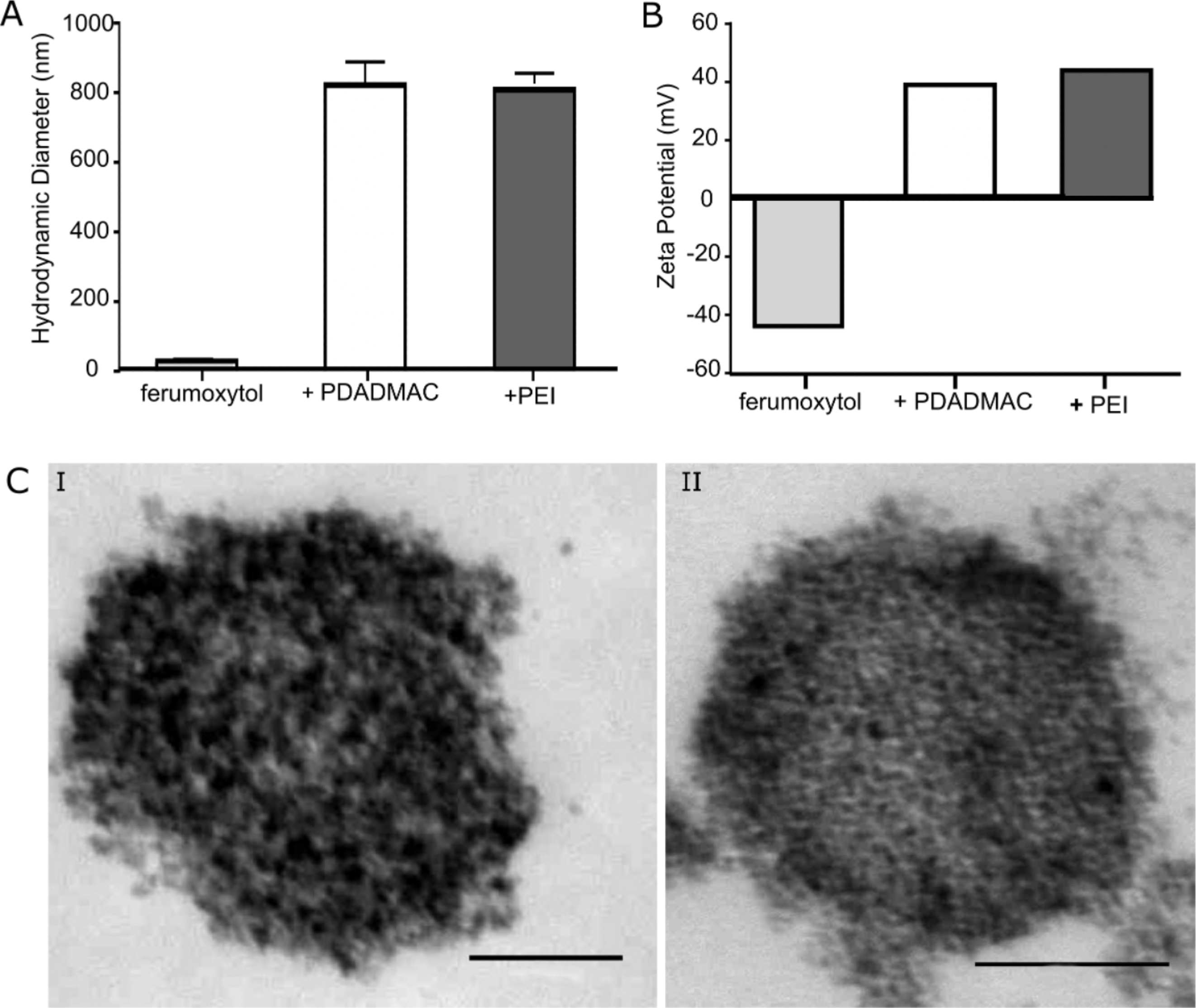
Characterization of uncoated, PDADMAC-, and PEI-coated ferumoxytol MNPs. (A) Hydrodynamic diameter of MNPs determined by DLS scans. (B) Surface charges (zeta potential) of MNPs obtained by Zetasizer measurements. (C) TEM images of (I) PDADMAC-coated ferumoxytol and (II) PEI-coated ferumoxytol clusters. Scale bars, 100 nm.

### Dendritic cell culture

Various DC subsets were differentiated from hematopoietic DC progenitors of bone marrow suspensions from C57BL/6 mice (Charles River; Sulzfeld, Germany) as previously described (Figure 1B) [21,22]. In brief, for the generation of the inflammatory type of DCs, Flt3+ DC progenitors were obtained from mouse bone marrow cells after 7 days of culture in RPMI 1640 medium supplemented with 10% fetal calf serum, 2 mM L-glutamine, 100 U/ml penicillin/streptomycin, and 50 μM β-mercaptoethanol (all from Life Technologies, Darmstadt, Germany) containing recombinant murine SCF (100 ng/ml), 25 ng/ml Flt3-ligand (Flt3L; PeproTech, Hamburg, Germany), 5 ng/ml hyper IL-6 (a kind gift from Dr. S. Rose-John, University of Kiel, Kiel, Germany), 40 ng/ml recombinant long-range IGF-1, 20 U/ml recombinant mouse GM-CSF, and 10^−6^ M dexamethasone (Sigma-Aldrich). After 7 days, differentiation of DC progenitors into inflammatory DCs was induced in culture medium supplemented with 200 U/ml recombinant murine GM-CSF [21].

A fraction of mouse bone marrow cells were cultivated for 7 days as described above to obtain steady-state cDCs and pDCs but in the absence of GM-CSF and dexamethasone [22]. After 7 days, such DC progenitors were differentiated into cDCs and pDCs using 50 ng/ml Flt3L only.

Cell numbers were determined with an electronic cell counting device (CASY1, Roche, Penzberg, Germany). Mice were maintained under specific pathogen-free conditions in the central animal facility of the RWTH University Hospital Aachen. All animal experiments were approved by local authorities in compliance with the German animal protection law and EU guidelines (2010/63/EU) for animal protection.

### Flow cytometry

Flow cytometry was used to analyze the DC phenotype and the effect of labeling as previously described [18]. Surface antigen expression on DC progenitors and DCs was examined using the following antibodies: Pacific Blue (PB)-conjugated anti-CD11b (clone M1/70), phycoerythrin-cyanin 7 (PE-Cy7) or fluorescein isothiocyanate (FITC)-conjugated anti-CD11c (N418), Allophycocyanin (APC)-conjugated anti-CD115 (AFS98), PE-Cy7-conjugated anti-CD117 (ACK), PE-conjugated anti-CD135 (A2F10), peridinin chlorophyll protein-cyanin 5.5. (PerCP-Cy5.5)-conjugated anti-B220 (RA3-6B2) APC-conjugated anti-MHC class II (MHC II; clone M5/114.15.2), and APC-conjugated anti-SiglecH (eBio440c) were purchased from eBioscience (San Diego, CA). PE-conjugated anti-CD24 (M1/69), FITC-conjugated anti-MHC II (2G9), and PerCP-Cy5.5-conjugated anti-Gr1 (RB6-8C5) were obtained from BD Biosciences (Heidelberg, Germany). Respective isotype controls were from BD Biosciences or eBioscience. Stained cells were analyzed with FACS Canto II flow cytometer (BD Bioscience), and data were evaluated using FlowJo software (TreeStar, Ashland, OR).

### MNP-labeling of DCs

Sterile solutions of ferumoxytol MNPs were sonicated for 30 min prior to cell labeling and used with a final iron concentration of 10 μg/ml. GM-DCs and FL-DCs were seeded at 2 × 10^6^ cells/ml and incubated with MNPs for 24 h. Cells were harvested and washed in a PBS solution containing 2% FCS before being subjected to magnetic separation (MACS; Miltenyi Biotech, Bergisch Gladbach, Germany) as previously reported [9,16]. The labeling efficiency was calculated after the magnetic separation of the retained cell (Equation 1). Cell numbers of retained and not-retained cells were determined using CASY1 cell counter.

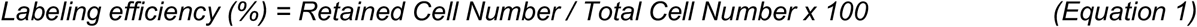

### Cell viability assessment after MNP-labeling

Cytotoxicity of PE-coated and uncoated MNPs on various DC subsets was evaluated using the Zombie Aqua Viability Kit (BioLegend, Fell, Germany) according to manufacturer’ s protocol. Briefly, 5 × 10^5^ cells after MNP-labeling were incubated in a 1:100 (v/v) dilution of the Zombie Aqua dye in a total volume of 50 μl PBS for 20 min at room temperature in the dark. After washing with PBS, cells were analyzed with the FACSCanto II flow cytometer and data were evaluated using FlowJo software.

### Quantification of cellular iron content

Intracellular iron concentration was determined using colorimetric ferrozine iron assay as previously described [9,16]. Briefly, 1 × 10^6^ cells were lysed in 100 μl 50 mM NaOH and ascorbic acid was used to reduce Fe^3+^ to Fe^2+^ ions that form a chelate complex with ferrozine (3-(2-pyridyl)-5,6-bis(phenyl sulfonic acid)-1,2,4-triazine; Sigma-Aldrich). The absorbance of Fe^2+^-ferrozine was measured at 550 nm using a FLUOStar OPTIMA plate reader (BMG Labtech, Ortenberg, Germany) and compared to the absorbance of FeCl_3_ standards.

### Prussian Blue staining of intracellular iron

1 × 10^5^ cells labeled with MNPs were centrifuged onto a glass slide using a cytospin centrifuge (Thermo Fisher Scientific, Dreieich, Germany) to visualize the total iron uptake of cells. After washing in ddH2O, glass slides were subjected to Prussian Blue staining using a 1:1 solution (v/v) of 10% K_4_[Fe(CN)_6_] (Sigma-Aldrich) and 20% HCl for 20 min. After washing in ddH2O, glass slides were counter-stained with Neutral Red dye (Roth, Karlsruhe, Germany) and mounted with coverslips using mounting medium (Dako, Hamburg, Germany). Sample images were obtained using the Leica DM6000B microscope and Diskus acquisition software (Hilgers, Koenigswinter, Germany).

### Transmission electron microscopy

Transmission electron microscopy (TEM) images were obtained from 1 × 10^6^ magnetically sorted DCs. Cells were fixed with 3% (w/v) glutaraldehyde and embedded in 2% agarose. Samples were stained with OsO_4_, embedded in Epon and cut into 70 nm thick slices. Samples were analyzed without further contrast enhancement using a Philips EM 400T electron microscope at 60 kV equipped with a CCD camera (MORADA, Olympus, Hamburg, Germany). TEM images were analyzed using NIH ImageJ software.

### Statistical analysis

Numerical data were analyzed for significance by one-tailed Student’s *t* test with GraphPad Prism software (GraphPad, La Jolla, CA). A *p*-value below 0.05 was considered significant.

## Results

### Characterization of PE-coated ferumoxytol MNPs

MNPs for biomedical applications generally possess a core-shell structure, whereby the shell surface is a crucial factor to impart good colloidal stability and biocompatibility, and ideally provides a scaffold for further functionalization. Ferumoxytol is a colloidal stable nanoparticle formulation with a core size of 6 nm and a polyglucose sorbitol carboxymethylether shell. We determined for ferumoxytol a hydrodynamic diameter of 29 nm and a ζ-potential of - 44 mV (Figure 1A and B). We frequently use layer-by-layer (LbL) assembly of high and low molecular weight (MW) polyelectrolytes around MNPs to improve cellular responses, including uptake, intracellular localization and processing of MNPs, and thus on MRI properties of labeled cells [16,18].

Here, based on previously established conditions for PE-coating of ferumoxytol, we used positively charged low MW polydiallyldimethylammonium chlorid (PDADMAC) and high MW polyethyleneimine (PEI) for coating (Figure S1A). LbL assembly of PDADMAC and PEI on ferumoxytol resulted in MNPs with the increased hydrodynamic diameter and positive surface charge, indicating successful PE coating (Figure 1A and B). TEM imaging revealed that ferumoxytol MNPs aggregated upon PE-coating to larger size clusters (Figure 1C). Under the chosen conditions for coating, the low MW PDADMAC-coated ferumoxytol MNPs (referred in the following as + PDADMAC) and high MW PEI-coated ferumoxytol particles (mentioned in the following as + PEI) were most similar in respect to surface charge, hydrodynamic diameter, and cluster size (Figure 1) and were therefore selected for further cell labeling studies.

### Differentiation of steady-state and inflammatory DC subsets

A number of protocols have been developed for generating DCs *in vitro* from bone marrow or cord blood-derived HSC or blood monocytes. Our group has successfully established two-step culture systems that allow the amplification of fms related tyrosine kinase 3 positive (Flt3+) DC progenitor populations in the first step under growth-promoting conditions, which in a second step can be differentiated into fully functional DCs and thus yield high cell numbers [21,22]. Dependent on the amplification and differentiation condition that contains or lacks GM-CSF different DC subsets, including inflammatory DCs, as well as cDCs and pDCs, were generated.

Expression of the surface integrin CD11c represents a hallmark of fully differentiated DCs, whereas Flt3+ progenitors are CD11c negative but express the integrin CD11b (Figure 2 and Figure S1A). After 7 days under proliferating conditions, the differentiation of Flt3+ progenitors towards DCs was induced (referred to as day 0 in Figure 2A and Figure S1A) using either GM-CSF or Flt3 ligand (Flt3L), respectively. The Flt3+ progenitor grown in the presence of GM-CSF and dexamethasone readily differentiates within 6 days into fully functional DCs following administration of GM-CSF only (hereinafter referred to as GM-DCs). The majority of GM-DCs displayed a low to an intermediate expression of Major Histocompatibility Complex class II (MHC II) molecules on the cell surface (Figure 2A), which was indicative for an immature developmental stage of GM-DCs [21].

**Figure 2.**
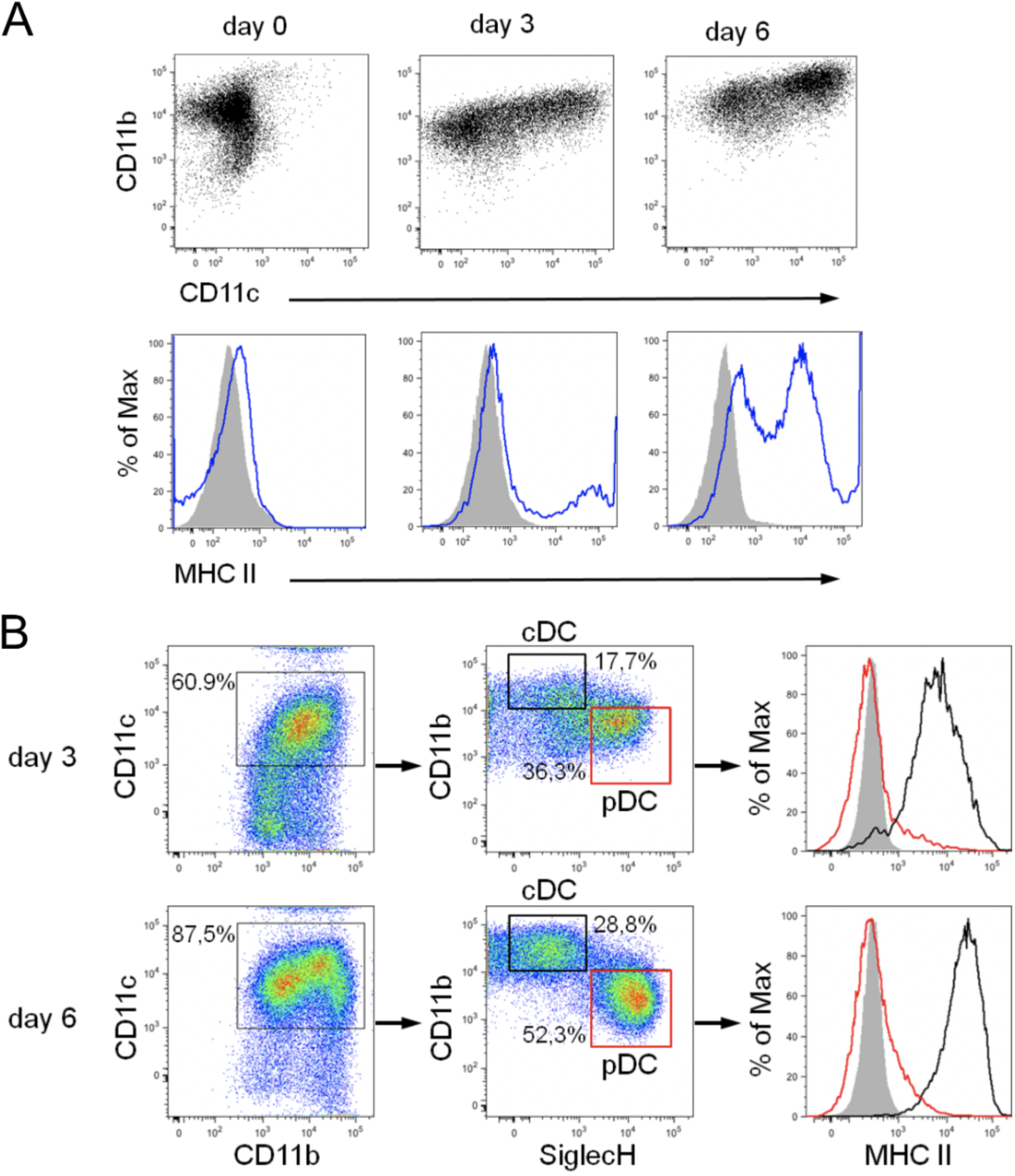
Phenotypic characterization of developing GM-DCs and FL-DCs during differentiation from Flt+ DC progenitors. (A) GM-DC development was monitored for CD11b and CD11c expression (upper panel), and MHC II expression (blue lines; lower panel) on days 0, 3 and 6 of differentiation using GM-CSF. (B) FL-DC development towards cDCs and pDCs on days 3 and 6 of differentiation using Flt3L was examined by gating on CD11c, CD11b, and SiglecH (left and middle panel). According to CD11b and SiglecH expression cDC and pDC populations were identified and MHC II expression within specific DC subsets is shown in the right panel (pDCs, red lines and boxes; cDCs, black lines and boxes). Gray histograms show isotype control staining. Representative flow cytometry data from 3-5 independent experiments are shown.

A modified two-step protocol was used that lacks GM-CSF and dexamethasone to obtain steady-state cDCs and pDCs [22]. This approach recapitulated more closely the *in vivo* DC development from a multipotent hematopoietic progenitor (MPP) through a common DC-restricted progenitor (CDP) intermediate stage [23]. Accordingly, under growing conditions, the DC progenitor populations were more heterogeneous reflected by different surface marker profiles: MPPs express high levels of c-kit (CD117) but are low to negative for Flt3 (CD135), while CDPs express c-kit at low levels and Flt3 at high levels (Figure S1A). CDPs also express CD11b and the macrophage-colony stimulating factor receptor (M-CSFR, CD115) at higher levels than MPPs (Figure S1A). Collectively, the *in vitro* amplified CDPs resembled a phenotype similar to their *in vivo* counterparts [24,25]. The modified growing conditions allowed differentiation of Flt3+ DC progenitors using Flt3L only that resulted in concurrent differentiation of both cDCs and pDCs within 6 days (Figure 2B and Figure S1A and B). Differentiated cDCs and pDCs exhibited an immature phenotype that resembles steady-state DC subsets in lymphoid tissues, such as spleen or lymph nodes [22]. In the bulk culture, we refer to both DC subsets in the following as FL-DCs. Mouse pDCs expressed CD11c, SiglecH, and B220 and downregulated CD11b expression, while cDCs expressed CD11c, CD11b and were negative for SiglecH and B220 (Figure 2B and Figure S1A, B). Moreover, both steady-state DC populations displayed a clearly distinct MHC II expression profile with definite MHC II expression in cDCs and the absence of MHC II expression in pDCs (Figure 2B) that is in line with the surface marker profile of their *in vivo* equivalents [26].

### Labeling of DCs with uncoated and PE-coated ferumoxytol

Next, MNP interaction with inflammatory and steady-state DCs was investigated. After incubation of cells with uncoated and PE-coated ferumoxytol particles for 24 h, the cells were subjected to magnetic sorting to separate MNP-labeled DCs and unlabeled DCs and thus to quantify the efficiency of MNP uptake. The labeling efficiency was calculated with the formula given in Equation 1, and the results are shown in Figure 3A. Labeling efficiency by uncoated ferumoxytol and PE-coated MNPs was found highly variable in GM-DCs and FL-DCs (Figure 3A). Uncoated ferumoxytol resulted in the labeling of only ~20% of FL-DCs in agreement with previously reported poor labeling efficiency [15,16]. In contrast, GM-DCs showed significantly higher labeling efficiency by uncoated ferumoxytol than FL-DCs. The PE-coating of MNPs resulted in increased cell labeling in line with previous studies but with different impact [16,27]. For FL-DCs, the MNP coating with both PEI and PDADMAC resulted in a four-fold higher labeling of FL-DCs than with uncoated MNPs (Figure 3A). In contrast, in GM-DCs, only the MNP-coating using PDADMAC improved MNP uptake by the cells, while PEI-coated MNPs showed no significantly improved labeling compared to uncoated MNPs. These results indicate that the improved labeling of GM-DCs by PDADMAC-coated ferumoxytol cannot be attributed solely to the positive surface charge. However, charge reversal by the PE-coating might indeed represent the primary cause for the improved uptake of MNP by FL-DCs. Taken together, the data suggest that different uptake mechanisms for MNPs are active in inflammatory and steady-state DCs, respectively.

**Figure 3.**
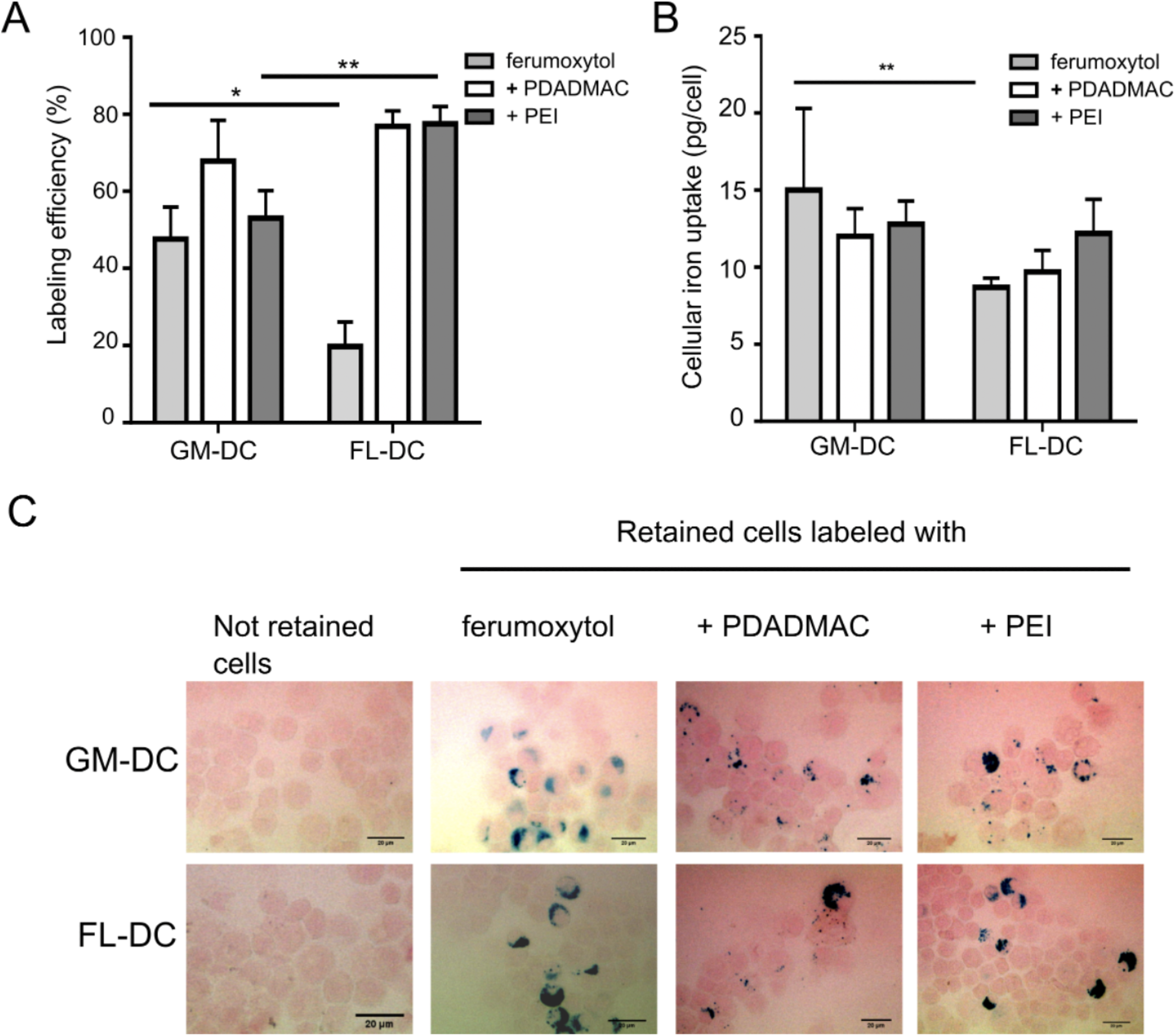
Uptake of uncoated and PE-coated ferumoxytol particles into DCs. Inflammatory DCs (GM-DCs) and steady state DCs (FL-DCs) were incubated with the respective MNPs for 24 h and subjected to magnetic separation of MNP-loaded cells from unlabeled cells. (A) Labeling efficiency was determined from numbers of retained and not-retained DCs after magnetic separation. (B) Intracellular iron concentration (pg/cell) of retained DC fractions after magnetic separation. Results in (A) and (B) are mean values ± SD (n=3; *: p<0.05; **: p<0.01). (C) Prussian blue staining of cytospins of magnetically retained and not-retained cells for detection iron deposits. Neutral red dye was used as counterstain. Scale bars, 20 μm.

The different MNP surface coatings and/or uptake mechanisms may impact on the intracellular loading of MNPs. To address this, we quantified the cellular iron content upon uptake of MNPs using a ferrozine-based colorimetric assay (Figure 3B) and visualized the MNP loading by Prussian blue staining (Figure 3C). It was found that GM-DCs were capable of taking up 15 ± 5.3 pg iron/cell with uncoated ferumoxytol particles, 12 ± 1.8 pg iron/cell with PDADMAC-coated MNPs and 12.8 ± 1.5 pg iron/cell with PEI-coated MNPs. The iron concentrations in FL-DCs were 8.7 ± 0.6 pg iron/cell with uncoated ferumoxytol particles, 9.7 ± 1.4 pg iron/cell with PDADMAC-coated ferumoxytol particles and 12.2 ± 2.2 pg iron/cell with PEI-coated ferumoxytol particles. No significant differences in the amount of intracellular iron were observed between GM-DCs and FL-DCs when PE-coated ferumoxytol particles were used. However, the internalized iron content was significantly higher in GM-DCs than in FL-DCs when labeled with uncoated ferumoxytol (Figure 3B). Notably, the increased labeling efficiency of FL-DCs with PE-coated MNPs was not reflected by elevated iron concentrations indicating a constraint in the uptake capacity.

The ascertained intracellular iron concentrations represent average values over all magnetically separated and hence labeled cells. Thus, the intracellular iron load was additionally visualized using Prussian blue staining to confirm the uptake of MNPs in labeled cells (Figure 3C). Iron deposits stained by Prussian blue showed a variable distribution of internalized MNPs in retained cells and revealed differentially sized agglomerates of the different MNPs within cells (Figure 3C). Furthermore, we did not observe any staining of iron in not-retained cells, confirming the absence of MNPs.

Labeling of steady-state DCs, including both cDCs and pDCs subpopulations and selection of labeled cells by magnetic sorting, was performed in bulk cultures. However, the analysis of cells after sorting by flow cytometry allowed discriminating labeled cDCs from labeled pDCs and to further address DC population-specific cellular responses to MNP uptake. Both cDC and pDC populations were identified according to differential CD11b and SiglecH expression (Figure 2B and Figure 4A). After 6 days of differentiation, around 30% of CD11c+ cells were identified as cDCs, and 50% were pDCs, while the other CD11c+ cells could not be clearly assigned to a specific subset (Figure 2B). A comparable ratio of 27.5% ± 1.8% (mean value ± SEM; n=3) cDCs and 49.0% ± 9.0% pDCs was also observed after further 24 h of culture (Figure 4A, B). The composition of FL-DCs after MNP-labeling for 24 h and magnetic separation was assessed accordingly (Figure 4A, B). After labeling with uncoated ferumoxytol, the ratio of cDCs and pDCs changed towards a higher proportion of labeled cDCs (52.6% ± 12.1% vs. 34.8% ± 10% of labeled pDCs) that further points to the poor labeling efficiency of uncoated ferumoxytol especially of pDCs (Figure 4C). Labeling with PDADMAC-coated MNPs was found equally efficient in both cDCs and pDCs as displayed by similar ratios as in unlabeled controls (25.8% ± 2.5% cDCs and 55.5% ± 5.1% pDCs). Labeling with PEI-coated MNPs resulted in a slight shift in the ratio of cDCs and pDCs towards labeled pDCs, demonstrating that PEI-coated MNPs are most effective for labeling of pDCs (Figure 4C).

**Figure 4.**
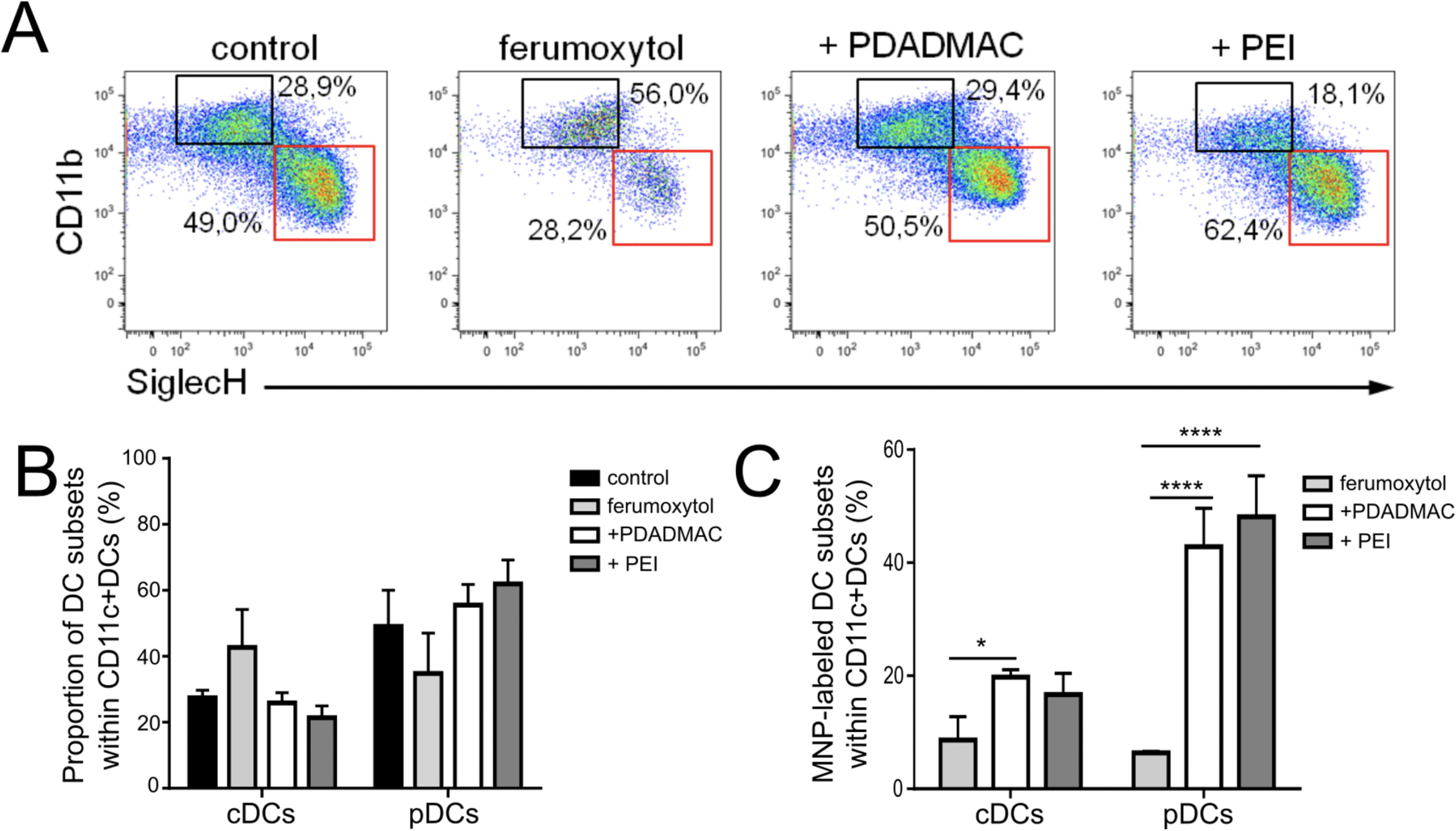
Uptake of uncoated and PE-coated ferumoxytol particles into steady state DCs. FL-DCs after uptake of respective MNPs and magnetic separation of MNP-loaded cells were analyzed by flow cytometry. Unlabeled cells were used as control. (A) CD11c+ cells were gated for CD11b and SiglecH as in Figure 2B to specify cDC (black boxes) and pDC (red boxes) subsets. Representative dot plots from 3 independent experiments are shown. (B) Proportion of DC subsets within MNP-labeled FL-DCs compared to unlabeled control FL-DCs. (C) Percentage of MNP-labeled DC subsets within all FL-DCs. Data in (B) and (C) are mean values ± SEM (n=3; *p<0.05,****p<0.0001).

#### Cytotoxicity assessment of uncoated and PE-coated MNPs upon uptake into DCs

We next examined whether MNPs affect cell viability of DCs after MNP-uptake by subjecting MNP-labeled and unlabeled cells to Zombie Aqua staining for detection of dead cells using flow cytometry (Figure 5). Unlabeled negative control GM-DCs showed a contingent of 5.2% of dead cells (Figure 5A). Unlabeled GM-DCs were stimulated with the bacterial wall component lipopolysaccharide (LPS) that leads to prominent DC activation [18] and 24 hours after LPS treatment, 9.6% of GM-DCs was found dead. After labeling GM-DCs with MNPs for 24 h and magnetic sorting the proportion of dead cells detected was 4.8% for labeling with uncoated ferumoxytol, 5.1% for labeling with PDADMAC-coated ferumoxytol, and 4.6 % with PEI-coated ferumoxytol. Thus, labeling of GM-DCs using uncoated and PE-coated ferumoxytol particles did not result in adverse cytotoxic effects at the concentration used here for labeling. Remarkably, particle size and surface charge also revealed no impact on cytotoxicity in GM-DCs in line with results from our previous studies [16,18].

**Figure 5.**
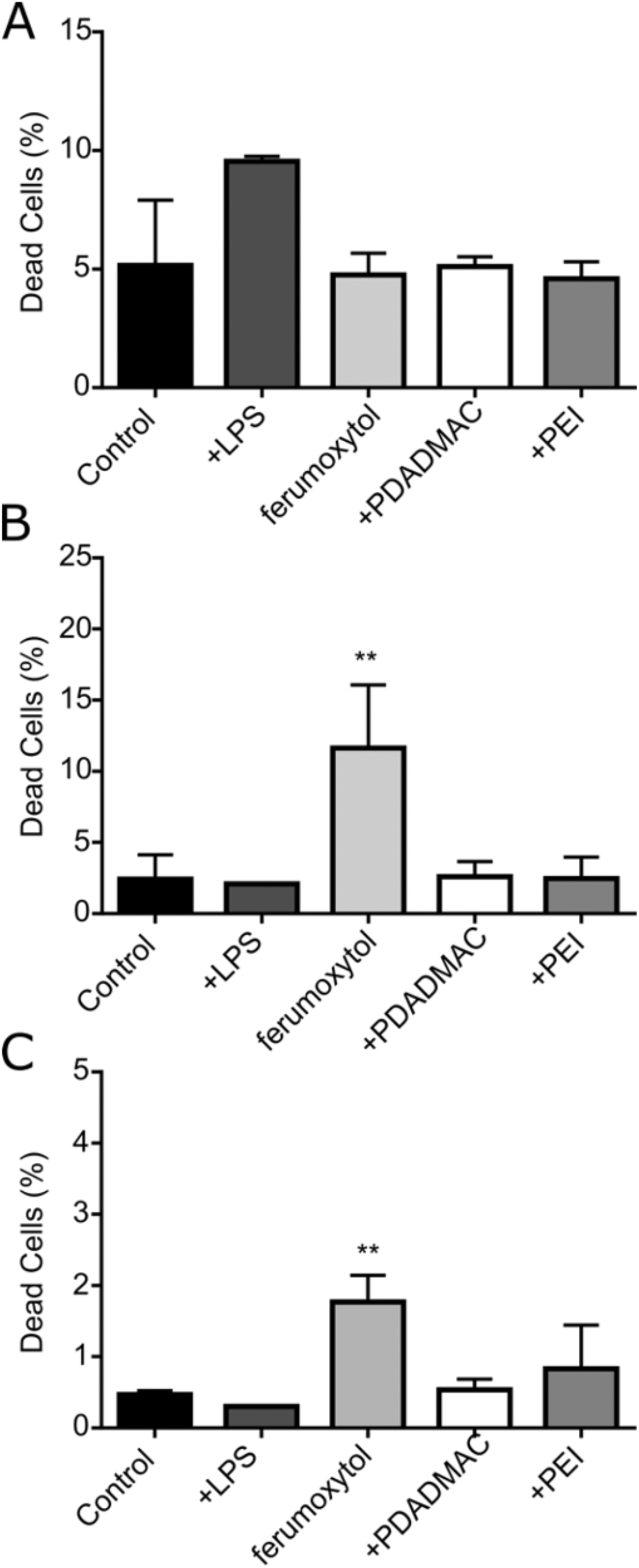
Biocompatibility of uncoated and PE-coated ferumoxytol in inflammatory and steady state DCs. Cellular toxicity was analyzed 24 h after MNP labeling and magnetic sorting using Zombie Aqua staining and analysis by flow cytometry. Bar diagrams show percent of dead cells in (A) GM-DCs, (B) FL-DC derived cDCs, and (C) FL-DC derived pDCs. Mean values ± SD (n=3) are plotted (** p<0.01).

In contrast to inflammatory GM-DCs, the response of steady-state FL-DCs to MNP labeling revealed a differential impact of uncoated and PE-coated ferumoxytol on cell viability after labeling (Figure 5B and C). Notably, the level of dead cells detectable in untreated and LPS stimulated FL-DCs was found lower compared to GM-DCs. In untreated cDC and pDC populations, 2.5% and 0.5% of cells, respectively, were determined as dead cells. Furthermore, stimulation of FL-DCs with LPS, unlike GM-DCs, did not result in an increased number of dead cells, although it led to effective DC activation (see below 3.5. and Figure 6C). However, the labeling of FL-DCs with uncoated ferumoxytol resulted in an approximately 3- to 5-fold increase in the number of dead cells in cDCs (11.6% ± 4.4%, mean value ± SD; n=3) and pDCs (1.7% ± 0.4%) yet on an overall low level. In contrast, the proportion of dead cDCs and pDCs after labeling with PDADMAC-coated MNPs (2.6% and 0.5%, respectively) or PEI-coated MNPs (2.5% and 0.8%, respectively) were not significantly changed compared to untreated FL-DCs. Thus, the PE-coating abolished the harmful impact of ferumoxytol in cDCs and pDCs, while yielding higher labeling rates and intracellular iron concentrations in these steady-state DCs.

**Figure 6.**
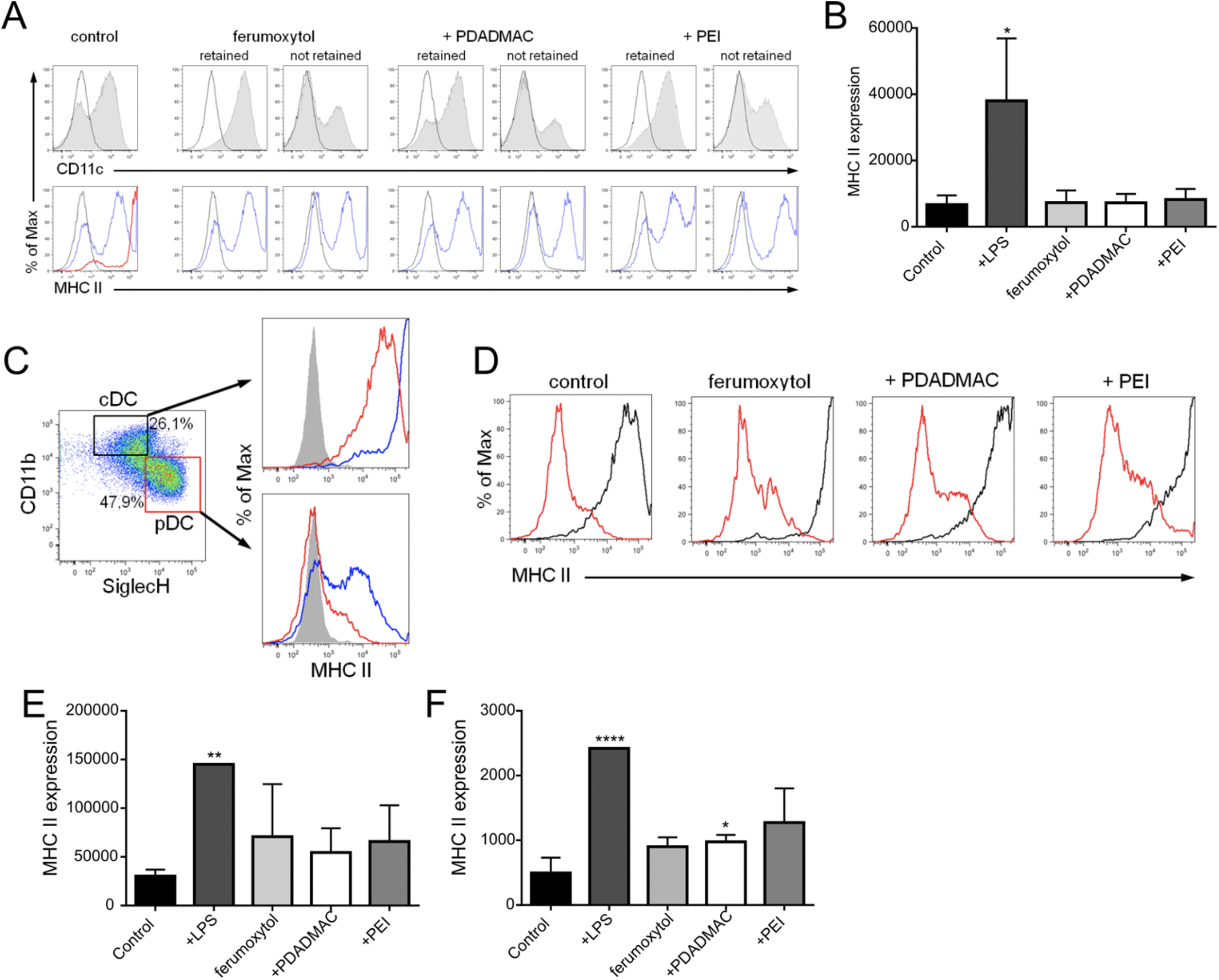
Immunophenotype of inflammatory and steady state DCs upon MNP uptake. MHC II expression as a measure of DC activation was analyzed 24 h after MNP labeling and magnetic sorting by flow cytometry. Unlabeled cells were used as controls displaying an immature phenotype. LPS stimulation of cells for 24 h was used as a control for DC activation. Representative data from at least 3 independent experiments are shown. (A) Surface antigen expression of CD11c (gray histograms) and MHC II (open blue histograms) on MNP-labeled GM-DCs. Activation of unlabeled DCs with LPS is depicted in red. Open black histograms show isotype control staining. (B) Bar diagram shows geometric mean of fluorescence intensity (geoMFI) values for MHC II expression ± SD on GM-DCs. (C-F) MHC II expression on MNP-labeled cDCs and pDCs. (C) DC subsets were identified according to CD11b and SiglecH expression and the MHC II expression (open red histograms) determined within specific DC subsets as before (see Fig. 2B). Gray histograms show isotype control staining. Activation of unlabeled cDCs and pDCs by LPS is depicted in blue. (D) MHC II expression on unlabeled and MNP-labeled cDCs (open black histograms) and pDCs (open red histograms). Results of MHC II expression on cDCs (E) and pDCs (F) are shown as geoMFI values ± SD.

### Immunophenotypic differences in steady-state and inflammatory DCs upon MNP-labeling

According to their specific antigen-uptake and -presentation function, DCs are well equipped with pattern recognition receptors that allow them to sense and uptake a wide range of antigens [28]. DC activation in response to foreign antigens leads to the maturation of DCs from their immature state that includes upregulation of MHC II on the cell surface required for effective antigen presentation to T cells. Here, we now assessed the immune stimulatory capacity of uncoated and PE-coated ferumoxytol particles upon uptake into both steady-state and inflammatory DCs, respectively. To this end, GM-DCs and FL-DCs were analyzed for MHC II expression by flow cytometry upon exposure to MNPs for 24 h and magnetic separation (Figure 6). As a control for effective DC activation, DCs were stimulated with LPS for 24 h. In the absence of MNPs, GM-DCs displayed low to intermediate expression levels of MHC II, which increased to high levels after stimulation with LPS (Figure 6A, B). Importantly, MNP-labeled GM-DCs showed MHC II expression profiles similar to profiles of untreated cells, indicating that exposure to and uptake of MNPs did not lead to activation of inflammatory DCs (Figure 6A, B) in line with our previous results [18]. In addition, the comparison of MNP-labeled with non-labeled cells (i.e., cells not magnetically retained) revealed enrichment of not terminally differentiated DCs (i.e., negative for CD11c and MHC II) in the not-retained fraction concurrent with the enhancement of terminally differentiated DCs in the MNP-labeled fraction (Figure 6A) that highlights the selective uptake capacity of DCs for MNPs.

In contrast to GM-DCs, the steady-state FL-DCs revealed a different MHC II expression profile with a more homogeneous expression of MHC II in cDCs at intermediate levels and the absence of MHC II expression in pDCs (Figure 3B and Figure 7C). LPS stimulation of FL-DCs showed a differential impact on FL-DC activation that led to the maturation of the entire cDC population to high-level MHC II expressing cells, while for pDCs, only a fraction of cells respond to LPS by upregulation of MHC II to intermediate levels (Figure 7C). Remarkably, and in contrast to inflammatory DCs, uptake of MNPs induced upregulation of surface MHC II expression in both cDCs and pDCs, although to a lesser extent than LPS (Figure 7D-F). Taken together, the results indicate that sensing and response to MNPs are differentially controlled in inflammatory DCs and steady-state cDCs and pDCs.

## Discussion

Surface coating of MNPs using polyelectrolytes possess great potential to tailor MNP properties for their use in various biomedical applications, including cell labeling and tracking by MRI. In this study, we demonstrated that specific PE coatings caused different cellular responses in distinct DC subpopulations. We found a selective labeling capacity of PE-coated MNPs dependent on the DC subtype together with a differential impact on the cytotoxic as well as immunomodulatory consequences of both uncoated and PE-coated ferumoxytol.

Engineered iron oxide MNPs are increasingly harnessed as advanced tools for medical applications, including cell labeling and tracking, targeted drug release, non-invasive monitoring of therapy, and vaccination [14,29]. Moreover, iron oxide-based MNPs are considered to be most suitable for combining a number of these applications into a single multifunctional formulation, thus providing an accurate nano theranostics tool.

Currently, ferumoxytol is the only FDA approved iron oxide-based MNP formulation initially launched for iron replacement therapy. It is recently investigated extensively as a contrast agent in MRI since it shows fewer side effects such as allergic or idiosyncratic reactions than other contrast agents [14]. Moreover, ferumoxytol does not entail a risk for the development of nephrogenic systemic fibrosis and thus may substitute gadolinium-based contrast agents as a blood pool agent in a number of MRI applications. Accordingly, attempts have been made to use ferumoxytol as a contrast agent for labeling and tracking of cells *in vivo* using MRI, including mesenchymal stromal cells, neural stem cells, and immune cells, including T cells, monocytes, and DCs [10,24,27,30]. The uptake by macrophages *in vivo* is being explored as a novel imaging approach for the assessment of lymph nodes, tumors, and vascular lesions. For example, in a preclinical model of autoimmune myocarditis, the iron oxide MNPs ingested by macrophages improved the distinction of areas of severe inflammation by MRI compared to conventional T2-weighted and gadolinium-enhanced MRI [31]. However, in a more recently published clinical study in patients with acute myocarditis, ferumoxytol-enhanced MRI was found unable to identify myocarditis by detection of macrophage activity [32]. As a possible explanation for the contradictory finding to the previous study, the authors discuss the limited uptake capacity of ferumoxytol by macrophages compared to other iron oxide MNPs. This is in line with previous studies that found no efficient labeling of cells with ferumoxytol alone or in combination with protamine [10,15]. However, a combination of ferumoxytol with protamine and heparin were described to result in improved labeling of neural stem cells, bone marrow stromal cell, monocytes, and T cells and an increase in T2 relaxivity compared to ferumoxytol alone [27].

We frequently use layer-by-layer (LbL) assembly of polyelectrolytes for coating of MNPs to improve cellular responses, including uptake, intracellular localization and processing of MNPs, and thus on MRI properties of labeled cells. For example, coating of oleate-stabilized MNPs with PDADMAC resulted in a more dense agglomeration of MNPs within endosomal compartments of DCs, resulting in a larger magnetic susceptibility effect (T2*) when compared with loosely packed MNPs that were coated with polystyrene sulphonate or chitosan [18]. Noteworthy, the observed differences in MRI contrast-agent properties of PE-coated MNPs were not related to the total amount of iron taken up by cells, as chitosan-coated MNPs yielded the highest iron concentration in DCs, but exhibited inferior performance in MRI [18]. Here, we used the positively charged polyelectrolytes PEI and PDADMAC for the coating of ferumoxytol. PEI and PDADMAC have proven to be excellently biocompatible as PE coating for MNPs and allowed stable colloidal coating of ferumoxytol [16,18]. Remarkably, the PE coating of ferumoxytol has significantly improved the labeling of steady-state DC by up to 4-fold. In contrast, uncoated ferumoxytol was already incorporated by ~ 50% of the inflammatory GM-DCs, and the PE coating did not result in substantially improved cell labeling.

Patient-derived GM-DCs are to date the prevailing DC subtype used in autologous cell-based immunotherapy studies [6]. Another approach, however, is to harness the available steady-state cDCs and pDCs *in vivo* by direct targeting of specific subsets and to activate their subset specific properties depending on the type of disease [3]. Accordingly, attempts are made by, e.g., using specific antibodies that recognize and bind to unique subsets as carriers for antigens, drugs or immune regulatory factors [33]. Besides, very similar approaches have emerged using nano carrier-based delivery systems, including iron oxide-based MNPs [34]. A wide range of nanoscale materials has been developed that can serve as platforms for assembling various antigens, adjuvants, and other immunomodulatory reagents bound to the surface of and/or enveloped in such nanocarriers. Interestingly, a number of such antigen and adjuvant factors represent polyelectrolytes that target specific immune pathways. One such pathway is activated by toll-like receptors (TLRs), and TLR agonists are intensively explored as molecular adjuvants for vaccination. This includes negatively charged nucleoside analogs that act as specific TLR agonists, such as double-stranded RNA or the synthetic analog poly(I:C) for TLR3, bacterial or viral single-stranded RNA for TLR7 or unmethylated DNA oligonucleotides (ODN) containing CpG motifs for TLR9. For example, simultaneously applied poly(γ-glutamic acid)-based NPs loaded with a tumor model antigen (OVA) or with poly(I:C) induced higher anti-tumor activity as compared to the activity without NPs [35]. Even more elegantly, the group of Jewell et al. [36] used LbL-assembly of positively charged OVA peptide (SIINFEKL) and negatively charged poly(I:C) around calcium carbonate templates where the core was finally removed using a chelator to create hollow capsules. These immune polyelectrolyte multilayers (iPEMs) were able to activate steady-state DCs to a greater extent than dose-matched soluble factors alone [34].

The TLR specificity of agonists can be further exploited to target specific DC subsets since DC populations express non-overlapping sets of TLRs [28]. For example, in humans, TLR9 is expressed by pDCs but not inflammatory DCs, and thus CpG ODN acts particularly on pDCs to induce type I interferon (IFN) production.

The synthetic polyelectrolytes PEI and PDADMAC used in our study elicited a slight adjuvant effect in steady-state DCs similar to ferumoxytol alone but to a much lesser extent than the TLR4 agonist LPS. In contrast, all NP formulations were immunological inert in inflammatory DCs, suggesting that GM-DCs, unlike FL-DCs, lack the respective recognition receptors.

A hitherto unrecognized immunomodulatory activity of ferumoxytol has been described in a recent study on the mechanism of tumor growth inhibition by ferumoxytol [37]. While ferumoxytol showed no direct cytotoxic effects on tumor cells, the study by Zanganeh et al. [37] revealed that ferumoxytol expedited the recruitment of macrophages towards tumor cells and induced a phenotypic shift towards pro-inflammatory M1 polarization. Tumor-associated macrophages generally develop towards an M2-like phenotype at a later stage of tumor progression. At the same time, M1 polarization requires activation by canonical IRF/STAT signaling pathways activated by IFNs and TLR signaling [38]. Ferumoxytol then elicits the production of reactive oxygen species (ROS) via the Fenton reaction in M1 macrophages, thereby inducing the apoptosis of cancer cells [37]. It is tempting to speculate that ferumoxytol is recognized and acts via TLRs on macrophages, but the precise mechanism has not yet been addressed. According to our data, however, it appears unlikely that ferumoxytol is recognized by TLR4, which is highly expressed in macrophages as well as GM-DC. In our study, both PE coated and uncoated ferumoxytol did neither result in activation nor apoptosis induction of inflammatory DC, in contrast to stimulation with the TLR4 ligand LPS. In contrast, uncoated ferumoxytol increased apoptosis in steady state DCs probably by generation of ROS through the Fenton reaction. Thus, the differential impact of ferumoxytol on the cytotoxicity of GM-DCs and FL-DCs further points towards DC subset specific pathways activated in responses to the MNPs.

## Conclusion

LbL assembly of polyelectrolytes around MNPs represents a versatile means to tailor MNP surface properties. Here, we found that high MW PEI and low MW PDADMAC are well suitable for the coating of ferumoxytol to improve the cell-labeling efficacy of various DC subtypes. PE-coated ferumoxytol was essentially non-toxic to labeled cells. Moreover, PE coating diminished adverse cytotoxic effects of uncoated ferumoxytol in labeled steady-state DCs.

Polyelectrolyte coating of MNPs can be conceived to endow particle surfaces with desired properties. This can be employed, e.g. (i) for enhanced or even targeted uptake into cells, thereby augmenting their performance as contrast agents for MRI and (ii) to provide additional functionalities such as immunoregulatory activities. For clinical use, immunomodulatory activities provided by the MNP formulation may have beneficial effects, e.g., such as an adjuvant function in vaccination applications. However, the careful assessment of such immunomodulatory activities is inevitable to avoid deleterious side effects of MNPs beyond cytotoxicity. The specific antigen-uptake and -presentation, as well as immune regulatory function of distinct DC subpopulations, makes them particularly important for the study of such MNP properties. Depending on the molecular pattern of the MNP shell, specific DC subsets - as shown here - can be expected to recognize MNPs as potentially foreign antigens, therefore providing the capability to biologically sense MNP shell chemistry. This can lay the foundation for further functionalization of MNPs that will combine efficient labeling of DCs for cell tracking with additional activities that impact on DC function. In this respect, the combination of ferumoxytol coated with clinically approved polyelectrolytes should expedite a faster translation of this approach to clinical use.

## Acknowledgements

We would like to thank H. Koenigs, RWTH Aachen University Hospital, Department of Pathology, for assistance with electron microscopy. This work was supported by the Flow Cytometry Facility, a core facility of the Interdisciplinary Center for Clinical Research (IZKF) Aachen within the Faculty of Medicine at RWTH Aachen University. Part of this work was supported by the Excellence Initiative of the German federal and state governments (ERS OPSP006 to J. E. W. and T. H.) by RWTH Aachen University, Germany. T. H. received a grant from the START program of the Faculty of Medicine, RWTH Aachen University. N. C. thanks the CEMACUBE program (Erasmus Mundus, Action 1, Common European Master’s course in Biomedical Engineering) for providing the scholarship.

