## Supplemental material for "Polyelectrolyte Coating of Ferumoxytol Differentially Impacts on Labeling of Inflammatory and Steady-State Dendritic Cell Subtypes"

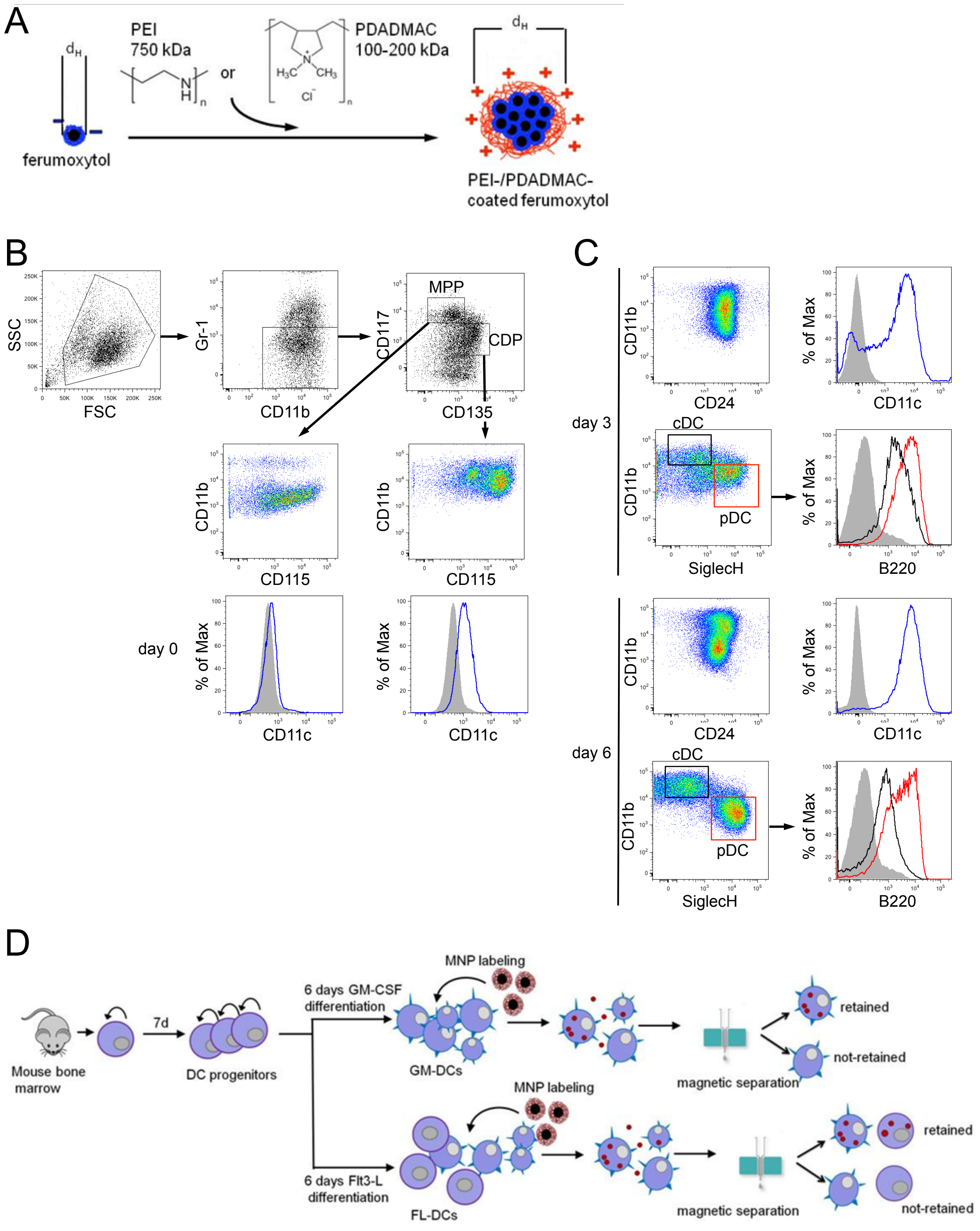


**Figure S1. DC labeling with PE-coated and uncoated ferumoxytol.** (A) Schematic representation of PE coating of ferumoxytol MNPs using PEI and PDADMAC; kDa, kilodalton; $d_H$, hydrodynamic diameter. (B-D) The two-step culture systems for differentiation of steady-state DCs (FL-DCs) and inflammatory DCs

(GM-DCs) from *in vitro* amplified Flt3+ DC progenitors and labeling with MNPs. (B) Phenotypic characterization of Flt+ DC progenitors for steady-state FL-DC differentiation towards cDCs and pDCs. Flt+ DC progenitors were examined by staining for Gr-1, CD11b, CD115, CD117/c-Kit, CD135/Flt3. According to CD117 and CD135 expression of Gr-1 negative cells, hematopoietic multipotent progenitor (MPP) and common DC-restricted progenitor (CDP) cells were identified (upper panel). Characteristic expression of CD11b, CD115 (middle panel), and CD11c (lower panel) in MPP and CDP are also shown. (C) FL-DC development towards cDCs and pDCs on days 3 and 6 of differentiation using Flt3L was examined by staining for CD11b, CD11c, CD24, SiglecH, and B220. According to CD11c, CD11b and SiglecH expression cDC and pDC populations were identified that display distinct pattern of expression for CD24 and B220 (pDCs, red lines and boxes; cDCs, black lines and boxes). Gray histograms show isotype control staining. Representative flow cytometry data from 3-5 independent experiments are shown. (D) Outline of MNP labeling and magnetic separation of cells developed in the two-step culture systems.
